# Land use influences on freshwater biodiversity

**DOI:** 10.1101/2025.09.08.674815

**Authors:** Minghua Shen, Roel van Klink, Wubing Xu, Alban Sagouis, Jonathan M. Chase

## Abstract

Land-use change is a major driver of biodiversity loss, but most generalizations about its effects come from terrestrial ecosystems. In freshwater systems, impacts are often indirect, driven by changes in the surrounding landscape that alter flow, chemistry, and connectivity. Using species-level abundance data from 163 studies across 6 biological groups, we examined how urban, agricultural, and forestry land uses affect freshwater biodiversity at local and regional scales. All three land-use types reduced species richness and altered relative abundance structure compared to natural vegetation, with urban systems showing the strongest and most consistent declines. Only urbanization was associated with biotic homogenization, particularly among abundant species. Biodiversity responses to land-use change also varied with latitude and taxonomic group: tropical assemblages tended to differentiate across sites, while homogenization was more common at higher latitudes. Macroinvertebrates and fish showed the strongest overall declines. These findings challenge assumptions of generality across ecosystems and highlight the need to explicitly incorporate freshwater biodiversity into global change research and policy.

## Introduction

Biodiversity is changing at rates unprecedented in human history, along with the functions and services biodiversity provides ^1-3^. However, much of what we know about biodiversity change comes from terrestrial systems, where decades of research have been synthesized to show widespread losses, typically linked to land-use changes ^4-6^. While these patterns have shaped conservation priorities and policy frameworks, they are based largely on data from the land ^7,8^.

Freshwater ecosystems have been notably underrepresented in these synthetic analyses, despite supporting a disproportionate share of global biodiversity and are highly sensitive to land-use change^2,9,10^. Unlike terrestrial habitats, where land use typically drives biodiversity loss through direct habitat conversion, freshwater impacts are often indirect—mediated by runoff, altered flow regimes, and habitat fragmentation from changes in the adjacent landscape^11-13^.

The directional and disconnected nature of freshwater systems also reduces the capacity of freshwater species to track suitable habitats under land-use changes, which in turn alters the freshwater community structure^14,15^. However, we still lack a clear understanding of how different land-use types reshape freshwater biodiversity across spatial scales and biological groups.

Understanding how land-use change reshapes patterns of biodiversity requires more than just counting species^16-18^. While species richness remains the most widely used measure to track biodiversity change, it overlooks shifts in species composition and dominance that can fundamentally alter community structure and function^19^. Abundance-sensitive metrics (e.g., Simpson diversity) capture these changes and often reveal contrasting patterns from richness alone^20,21^. Biodiversity and its change are also scale-dependent^17^. Spatial turnover, measured by beta diversity, captures how communities become more similar (homogenized) or different (differentiated) across landscapes in response to disturbance^22-24^. Different land uses may either flatten or amplify spatial variation, depending on intensity and context^25-28^.

While land-use effects on biodiversity have been widely studied, few syntheses have evaluated how these effects play out in freshwater ecosystems across spatial scales, metrics, and taxa. Existing meta-analyses often overlook scale dependence or don’t distinguish between richness and abundance-weighted diversity, limiting their ability to detect consistent patterns^29,30^. To address this, we used a recently published global database of species-level abundance data^31^ (FreshLanDiv), and compiled 163 studies from streams, rivers, and lakes across 33 countries. We compared assemblages in natural areas with those in landscapes dominated by urban, agricultural, or forestry land use—the three most frequent human pressures represented in the data. We quantified land-use effects on species richness and relative abundance structure at both local (α) and regional (γ) scales, and assessed spatial turnover (β-diversity) within each land-use type. We also examined whether these responses varied geographically and across biological groups. Our aim was to test whether patterns observed in terrestrial ecosystems extend to freshwater biodiversity, or whether they follow distinct, scale-dependent trajectories.

## Results

### Land-use effects on biodiversity across scales

At both α (local scale, sites level) and γ scales (regional scale, aggregated all sites within one land-use category), species richness was on average highest in freshwater habitats surrounded by natural vegetation with little land use, and lowest in urban ecosystems (Fig. S1). At the same time, the abundance of individuals in a given assemblage was lowest in natural vegetation but highest in urban ecosystems (Fig. S2, S3).

We compared the effect size between land-use categories (impacted vs reference) for two measures of diversity: species richness (S), which counts all species equally, and effective number of species given Simpson’s diversity (Spie), which gives greater weight to common species and reflects the dominance structure of communities. At the local scale, we observed similar magnitudes of decline in both species richness (S) and effective number of species (Spie), but the likelihood of decline differed between the two metrics. Urbanization and agriculture consistently reduced both metrics, whereas forestry led to a moderate decrease in S, but had no detectable effect on Spie (Fig. 1A). At the regional scale, urbanization had nearly two times higher impact on Spie than S, indicating stronger effects on dominant species. In agricultural landscapes, both metrics declined to a similar extent, and in forestry there were again weak declines in S but no consistent impact on Spie (Fig. 1B). For β-diversity—the spatial variation in community composition within land-use types—we found little change in richness-based turnover (β-S) across land uses. However, abundance-weighted turnover (β-Spie), which reflects spatial changes among dominant species, declined under urbanization, consistent with biotic homogenization, but remained stable in agricultural and forestry systems (Fig. 1C).

**Figure 1.**
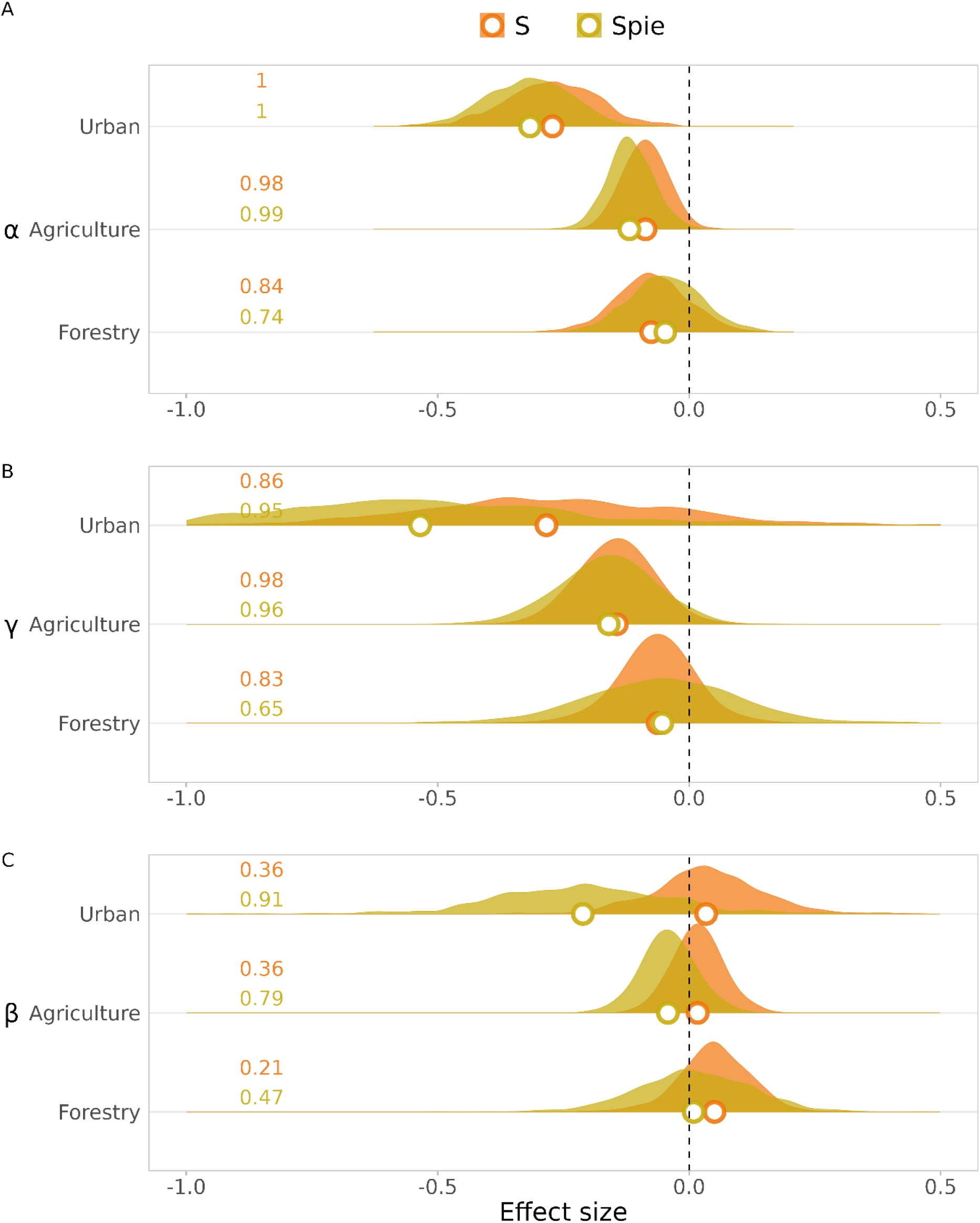
Density plot of the effect size (log-ratio) between urban, agriculture, forestry and natural vegetation across spatial scales. The vertical solid black lines show log-ratio equals to 0. The colored dots show the median value of the density plot of S (orange) and Spie (yellow). The colored numbers show the proportion (out of 1) of log-ratio below 0.

### Heterogeneity in responses across latitude, biological groups, and basin sizes

At the α scale, the negative effects of land use on both S and Spie were stronger at lower absolute latitudes (Fig.2 A, B), but no such variation was observed at the γ scale (Fig 2 C, D). As a consequence, β diversity, based on both S and Spie, declined with increasing absolute latitude. At lower latitudes (0–30°), land-use change led to stronger declines in the local-than regional-scale species richness, resulting in increased β-S and indicating biotic differentiation. In contrast, at higher latitudes (30–60°), β-S was declined, reflecting biotic homogenization (Fig.2E). However, β-Spie declined across all latitudes, indicating consistent homogenization driven by abundant species (Fig.2F).

**Figure 2.**
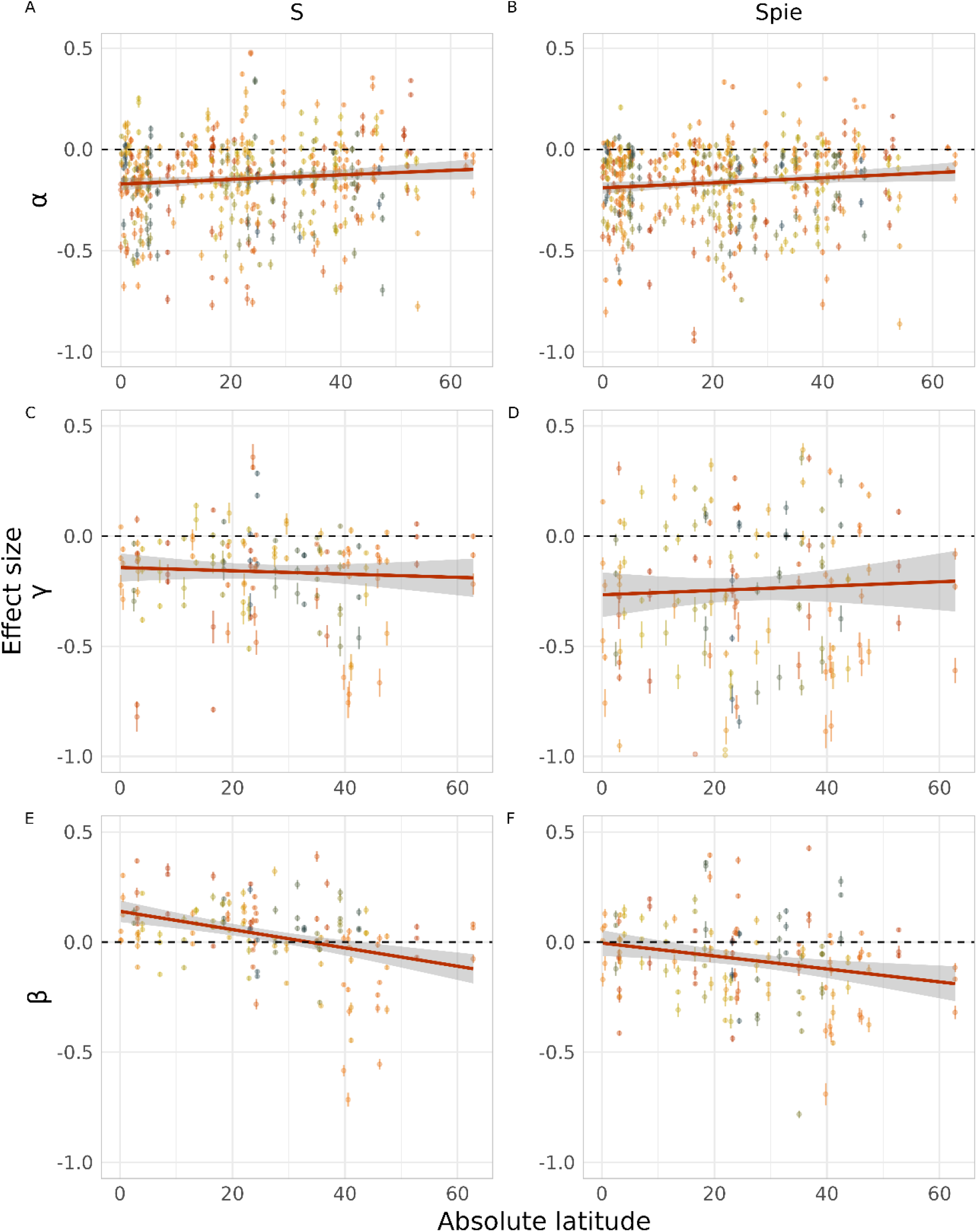
Relationship between absolute latitude and land-use effect sizes of α-S (A), α-Spie (B), γ-S (C), γ-Spie (D), β-S (E), β-Spie (F). The red solid lines represent the regression modeled using the “glm” function. The gray shadings show 95% credit intervals. Colored circles and vertical lines show the study-level estimate and 95% credible intervals. Different colors represent different studies.

For both macroinvertebrates and fish, the effect size of land use was negative for both diversity metrics at both α and γ scales. At the α scale, both S and Spie declined most in macroinvertebrates, followed by fish and algae. We observed a similar pattern at the γ scale for Spie (Fig.3 A, B), but γ-S declined most in algae, followed by macroinvertebrates and fish (Fig.3 C, D), suggesting stronger losses of rare species in algae communities. Land use had contrasting effects across groups for β diversity. For macroinvertebrates, β-S increased with land use intensification, indicating differentiation, whereas β-Spie decreased on average, suggesting homogenisation of abundant species. For fish, β-S did not change, but β-Spie decreased. In contrast, algae assemblages showed a decline in β-S, but no change in β-Spie (Fig.3 E, F).

**Figure 3.**
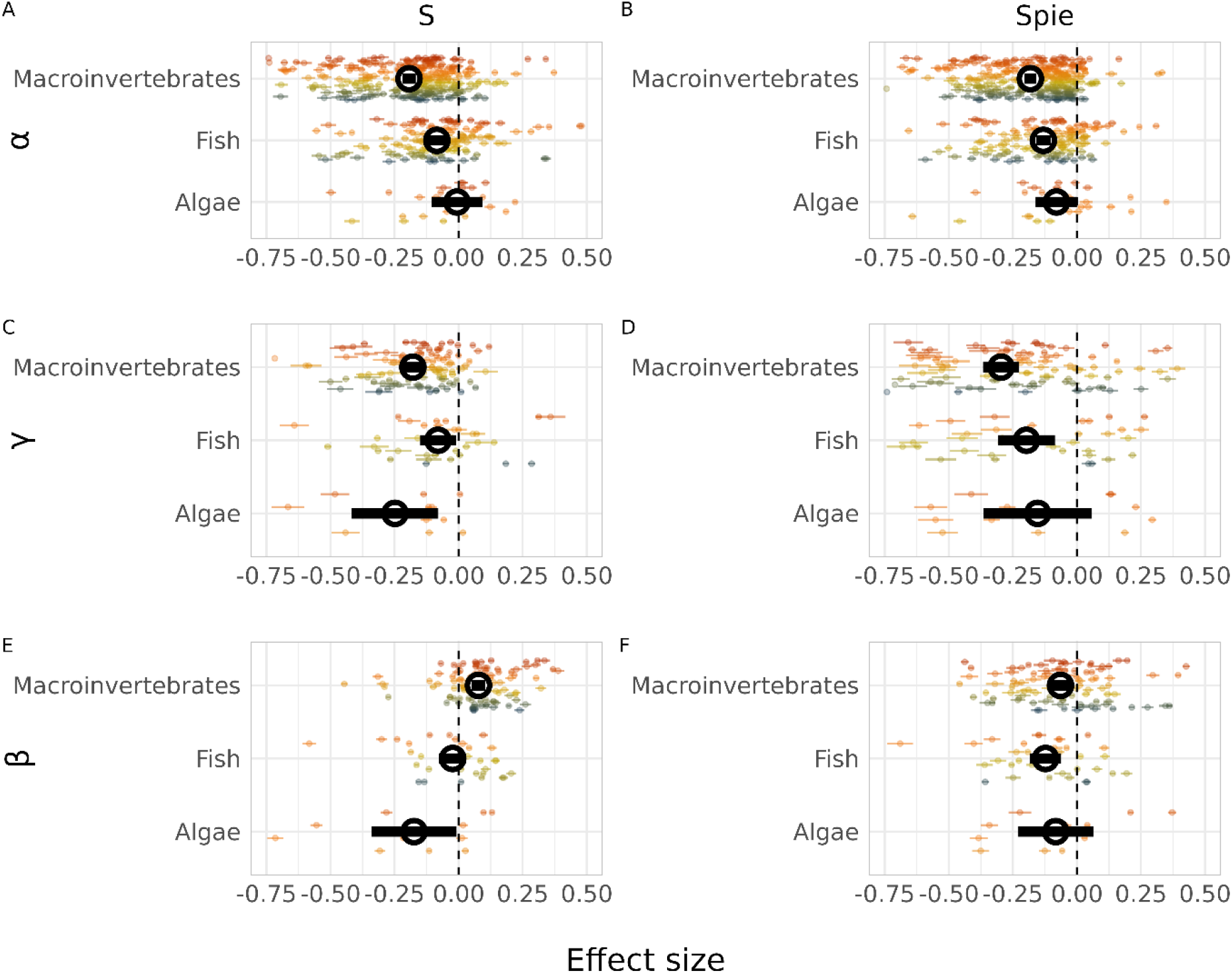
Study-level land-use effect sizes grouped by biological groups of α-S (A), α-Spie (B), γ-S (C), γ-Spie (D), β-S (E), β-Spie (F). The black circles show the average effect size for each biological group. The error bars show the 95% credible intervals. Colored circles and their error bars show the study-level estimate and the 95% credible intervals. Different colors represent different studies.

We did not find evidence that land-use effects varied among different basin sizes (Fig. S4).

## Discussion

Land-use change reshapes freshwater biodiversity, with consistent but context-dependent impacts on species richness, dominance structure, and spatial composition. Across 163 studies, we found reductions in species richness and abundance-weighted measures of diversity. These responses varied by spatial scale, geographic region, and taxonomic group.

Urbanization had the most pervasive effect, reducing species richness and disproportionately reducing abundance-weighted diversity (effective number of species based on Simpson index), particularly at larger scales. These patterns align with recent work highlighting the disproportionate loss of widespread, abundant species under strong human pressures^32,33^. This reflects not only species loss but the change in community dominance hierarchies through the disproportionate decline of locally common species. Such losses have also been observed in terrestrial birds and insects^34,35^ and likely stem from urban stressors such as impervious surfaces, altered hydrology, and contaminated runoff^36,37^. Urbanization consistently reduced both local and regional diversity. These effects were particularly strong for dominance-weighted metrics at the regional scale, leading to biotic homogenization through the loss of widespread, abundant taxa. Abundance-sensitive beta diversity declined, while incidence-based beta diversity changed little, reinforcing the conclusion that dominant species were especially affected. This erosion of community structure may impair ecosystem functioning and resilience^38,39^

In contrast, agricultural land use caused moderate but more uniform declines in both species richness and relative abundance structure, with minimal evidence for biotic homogenization. This contrasts with some reports of homogenization in agricultural water bodies^40^, and may reflect the greater heterogeneity of agricultural practices represented in our dataset—including variation in crop type, pastureland, and management intensity^41,42^. Such variation can preserve local environmental complexity and reduce compositional similarity across sites.

Forestry had the weakest overall effects. Local species richness declined slightly, but there were no consistent reductions in dominance-weighted diversity or beta diversity. This likely reflects the heterogeneity of forestry practices, which include deforestation, plantation forestry, and reforestation. These activities differ widely in their biodiversity impacts, and in some cases, reforestation and mixed-structure plantations may support greater habitat heterogeneity and biodiversity retention^43,44^.

Geographic context shaped these responses. In the tropics, local diversity declined more than regional diversity, suggesting loss of high-occupancy species and resulting in spatial differentiation^24^. These species may be specialists or abundance-limited taxa with narrow niches. In contrast, at higher latitudes, declines in both incidence-based and abundance-weighted beta diversity indicated homogenization driven by losses of both rare and dominant species. This is consistent with ecological patterns where tropical systems tend to support more specialized species with limited ranges and low local dominance^45,46^, while temperate systems host more generalists with broader ecological niches and larger population sizes^47-49^. Land-use change may favor a small set of generalists, driving homogenization especially at high latitudes.

Taxonomic patterns reveal further differences. Macroinvertebrates showed the strongest declines across all metrics, with reductions in both species richness and dominance structure, and declines in beta diversity. These shifts suggest replacement by a smaller pool of pollution-tolerant, widespread species^27,50,51^. Fish communities declined less steeply, possibly due to greater dispersal capacity or habitat buffering. Algal responses were more variable, partly due to limited data. These differences reflect trait-based sensitivities to land-use pressure, including dispersal ability, hydrological dependence, and tolerance to chemical stressors^44^.

While we leveraged the most comprehensive dataset currently available^31^, our ability to assess broad-scale regional effects was constrained by limited geographic metadata. Some regions, such as South America, were better represented due to intensive land-use transformation and active research networks^52,53^, while others such as southwest Australia and the Horn of Africa remain underrepresented due to limited research infrastructure^13^. Requiring site-level geographic data in order to spatially constrain the scale-dependent analyses excluded many otherwise relevant studies. However, our sensitivity analysis including studies that only collected local scale data confirmed that local diversity patterns remained consistent even when restricted to the subset of datasets that were also able to quantify regional scale changes (Fig. S5)

Our findings diverge from terrestrial syntheses, which report similar biodiversity losses under agriculture and urbanization and relatively consistent biotic homogenization across latitudes^5,6,54^. In freshwater ecosystems, urbanization emerges as the dominant driver of biodiversity loss, with stronger effects on common species and more pronounced latitudinal differences in homogenization. These differences likely arise from the physical structure and ecological dynamics of freshwater systems, including spatial isolation, limited dispersal, and vulnerability to surrounding land use. As biodiversity assessments advance^55,56^, integrating freshwater ecosystems is essential to understand global change and to design effective conservation strategies.

## Methods

### Data collection and selection

The data used in this analysis are derived from the FreshLandDiv database, which we created by aggregating data from existing studies on species abundance across different land-use categories. It includes all major freshwater biological groups: algae, macrophytes, zooplankton, macroinvertebrates, fish and amphibians. A complete description of the data collection and curation process is available in the datapaper^31^. This database contains eight land-use categories: natural vegetation, forestry, agriculture, urban, mining, impounded, unimpounded, and downstream of impoundment. From this database, we selected studies including natural vegetation as the reference status. We chose three most frequent human-impacted land-use categories: forestry, agriculture, and urban. We excluded dam related human impacts, including impounded, unimpounded and downstream, to focus on assessing land-use effects. The number of studies available for mining was low compared to the other three human-impacted land-use categories, so we excluded mining from the analysis to ensure the study numbers are comparable between land-use categories. If the original authors of a study provided several land-use categories, we chose the sites where the dominant land-use category occupied more than 50% and took the dominant land-use category as the land-use category for the site. These selections led to 163 studies with 3871 sites across 33 countries.

### Sampling standardization and calculation of biodiversity metrics

To assess the effects of land use on biodiversity, we examined whether these effects are scale-dependent and whether rare and abundant species respond differently. We selected two biodiversity metrics: species richness, which places high emphasis on rare species, and effective number of species given Simpson’s diversity, which incorporates the differences of relative abundance and is more sensitive to abundant species^16^. These two metrics were calculated across spatial scales (local alpha scale, and the regional gamma scale, as aggregate of all sites within a land-use category) to evaluate potential scale-dependent responses.

#### Sampling effort standardization

Prior to calculating biodiversity metrics at any scale, we ensured that sampling time, habitat type, methodology, and efforts were comparable between land-use categories.

Samples were organized hierarchically in a nested structure comprising sites, plots, and individual samples, with each site potentially containing multiple plots, and each plot comprising one or more samples. In studies where sites were sampled across different years or habitats, we subdivided sites into temporal or spatial blocks to ensure comparisons were made within the same study, year, and habitat. Before we calculate the metrics at any scale, we need to make sure the sample time, habitat, methods and sample efforts were comparable between land-uses. This allowed us to compare biodiversity between two land-use categories under consistent sampling conditions. For studies conducted in a single year and habitat, only one block was defined.

In addition to accounting for variation in sampling year and habitat, it was crucial to ensure that the sampling methods and efforts were comparable across different sites within the block. We have verified that the sampling methods were consistent within each block, while the sampling effort varies within some blocks, such as differing numbers of subsamples in spatial or temporal replicates. To address inconsistencies in sampling effort within a block, we standardized the data for studies where efforts were unequal. For example, if one site within a block had six samples and another had five, we resampled the site with six efforts 100 times to reduce its total abundance to 5/6 of its original. This produced 100 hypothetical communities, from which biodiversity metrics and abundances were calculated for each iteration. The final metrics were obtained by averaging these 100 values. This approach ensured robust comparisons between land-use categories.

#### Spatial extent standardization for γ and β diversities

According to species-area relationship, larger areas tend to support a greater number of species due to the availability of diverse habitats and resources^57^. For reliable comparisons of γ and β diversities, it is essential that the spatial extents across different land-use categories and blocks are comparable. Because some datasets have variation in spatial extent and/or the number of sites between land-use categories and blocks, we designed a process for creating comparable spatial extents in each study or block (we refer to this process as “cookie cutting”).

For the comparisons at the gamma and beta scales, we selected studies based on two criteria: (a) studies that sampled at least three sites spanning at least two land-use categories within each study and spatial block. (b) We prioritized studies where the land-use categories or spatial-temporal blocks had comparable areas (area difference ≤ ±50%), or where sites with comparable areas could be selected.

To standardize spatial extent, we employed two approaches:

1. Automated workflows: To retain as many sites as possible while ensuring spatial compatibility, we developed automated workflows for “cookie cutting” in studies with available geographic coordinates. When comparing land-use A and B, if land-use A had both the smallest number of sites (N_A_) and the smallest spatial extent (S_A_), the metrics of land-use A were calculated directly and cookie cutting was applied only to land-use B. We randomly selected 100 combinations of N_A_ sites from land-use B. For combinations with spatial extents within ±50% of S_A_, the metrics were calculated, and the mean across these combinations was used as the metric value for land-use B. In cases where land-use A had the smallest spatial extent (S_A_) but land-use B had the fewest sites (N_B_), cookie cutting was applied to both land uses. We randomly sampled N_B_ – 1 sites 100 times in each land use, retaining only combinations with spatial extents within ±50% of S_A_. The final metric for each land use was the mean of the accepted combinations. A detailed workflow is provided in Fig. S6.
2. Manual selection: For studies where spatial extent could not be standardised through automated workflows due to the absence of geographic coordinates, but where sitemaps were available, we manually selected sites to ensure comparability between land-use categories. We calculated polygon areas using WebPlotDigitizer (version 4.6) and manually selected sites, ensuring area differences remained within 50% of the minimum size (Figure S6). A total of 52 studies were included in the analysis after the selection process.

#### Calculation of metrics

We calculated three biodiversity metrics: the number of individuals (N), species richness (S), and the effective number of species given Simpson’s diversity (Spie)—at both α and γ scales, and two metrics (S, Spie) at the β scale, following the methodology outlined by Chase et al. (2018)^17^.

## Statistical analyses

We fitted Bayesian hierarchical models to investigate the relationship between biodiversity metrics and land-use. The response variables were biodiversity metrics, and the main fixed effect was land-use categories, a discrete variable representing different types of land-use. We did not include an intercept so that we could directly estimate the effects of each land-use type, since not all studies shared a common reference. We used random effects to allow the effect of land-use to vary at multiple levels in the nested structure of the dataset: study, block, site, and plot. This random effect structure controls for potential non-independence between observations within the same study, block, site, or plot. We assumed that most metrics across three spatial scales followed a lognormal distribution, except for S at α scale, which was assumed following a Poisson distribution to accommodate its integer values. In our analysis, α-S was the only metric with integers; γ-S contained non-integer values resulting from our spatial standardization (“cookie cut”) process.

For Bayesian inference and uncertainty estimates, we utilized the Hamiltonian Monte Carlo (HMC) sampler Stan^58^, implemented via the brms R package (version 2.20.4)^59^. Each model ran four chains of 2,000 iterations, with the first 1,000 iterations serving as warmup. We evaluated model fit using posterior predictive checks, confirming that the models effectively replicated the observed data (Supplementary Table 1 and Fig.S7).

We compared the fixed effect estimates between more-impacted land-uses and natural vegetation using posterior distributions to detect the effect sizes. For each land-use, we randomly drew 1,000 iterations of overall mean values to calculate the log ratio for each pair of land-use categories. There were three pairwise comparisons: forestry versus natural vegetation; agriculture versus natural vegetation; and urban versus natural vegetation.

To examine how effect sizes vary with latitude and across biological groups, we estimated study-level effect sizes by drawing 1,000 samples from the posterior distribution, comparing three more impacted land-use types to natural vegetation within each study. For latitudinal patterns, we assigned each study an average latitude based on the location of its sampling sites. If latitude data were missing, we used the geographic center of the corresponding province/state/country, depending on the most detailed available information. Bayesian hierarchical models were fitted to examine the relationship between absolute latitude and land use effect sizes. The effect sizes were used as the response variables, with absolute latitude as the fixed effect. Study was included as a random effect, allowing latitude effects to vary across studies. The model results can be found at Supplementary Table 2. To assess variation across biological groups, we categorized effect sizes by groups. For γ and β diversity, data were available only for macroinvertebrates, fish, and algae, so we restricted α diversity plots to these same groups.

We interpreted log ratios with 95% of distribution above or below zero as strong evidence for a directional trend. Moderate evidence was indicated by 90% to 95% of the distribution above or below zero, and weak evidence by 80% to 90%^35^.

### Sensitivity analysis

Most of the studies included in our synthesis were conducted in lotic ecosystems, such as rivers and streams. Given the large differences in relative basin sizes of the sites—from large rivers to small streams—the impact of land-use could vary depending on the size of the ecosystem. To explore this, we performed a sensitivity analysis using stream order as a proxy for ecosystem size. For this, we incorporated data from all studies with available geographical coordinates to determine the stream order for each site (2077 out of 3871 sites were included). We extracted stream order using the HydroRIVERS (version 1.0) dataset and conducted the same analysis as previously described. In this analysis, stream orders were treated as a random effect in the models. We then drew 1,000 samples from the posterior distribution for each study-level estimate and calculated the effect sizes between three more-impacted land-uses and natural vegetation. Finally, we examined the posterior distribution of the effect sizes across each stream order.

## Data availability

All the data used in this analysis are open access and available in the FreshLandDiv data paper: https://onlinelibrary.wiley.com/doi/10.1111/geb.13917. R code used for data standardizing and analysis are available on Github: https://github.com/chase-lab/Across_scale_analysis

## Acknowledgement

This work was supported by China Scholarship Council (Grant 202104910063), German Center for Integrative diversity Research (iDiv) Halle-Jena-Leipzig (DFG FZT 118-202548816), and ERC Advanced Grant (MetaChange) to J.M.C.. We thank the Biodiversity Synthesis group at iDiv for feedback.

## Author contributions

M.S., R.v.K., J.M.C. contributed to project design. M.S. compiled and analyzed the data with support from R.v.K., W.X., A.S., and J.M.C. M.S. wrote the first draft of the manuscript. All authors contributed to revisions.

