## Supplementary information for "Land use influences on freshwater biodiversity"

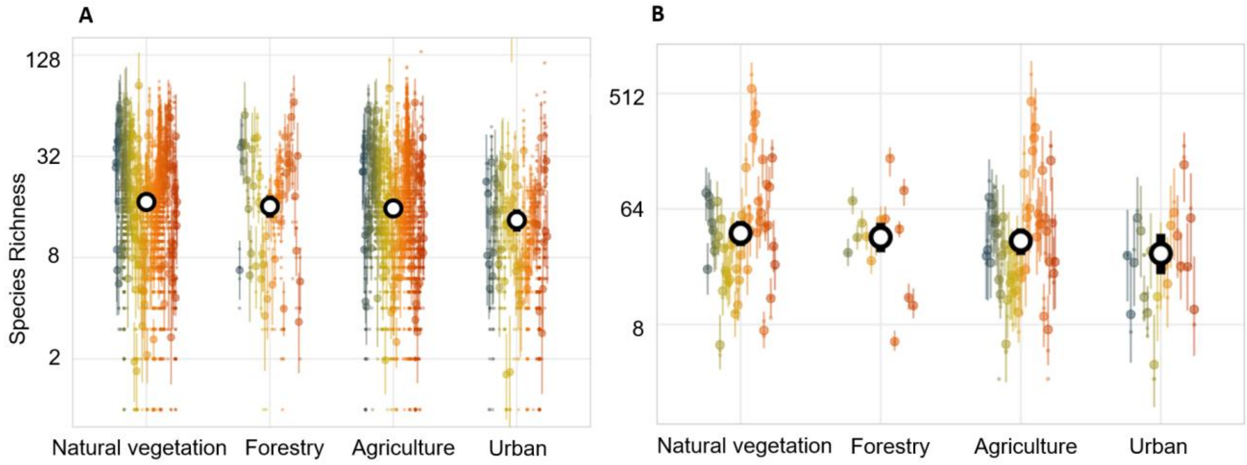


**Figure S1. Species richness across different land-use categories at α-scale (A) and γ-scale (B). Y-axes are on log-scales**. The black circles show the average richness of each land-use category. The black vertical lines show 95% credit intervals. Colored circles and vertical lines show the study-level estimate and 95% credible intervals; smallest colored dots show the raw data. Different colors represent different studies.

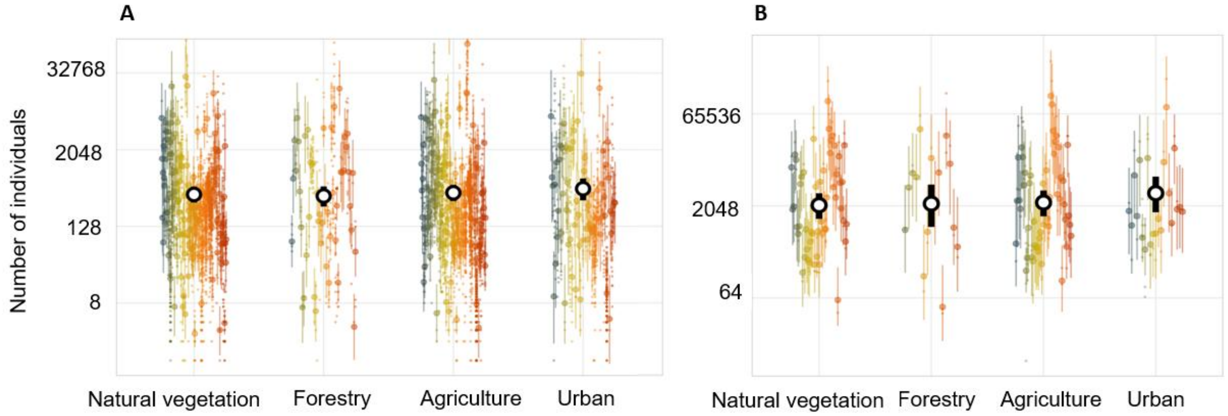


**Figure S2. Number of individuals across different land-use categories at (A) α-scale and (B) γ-scale.** Y-axes are on log-scales. The black circles show the average richness of each land-use category. The black vertical lines show 95% credit intervals. Colored circles and vertical lines show the study-level estimate and 95% credible intervals; smallest colored dots show the raw data. Different colors represent different studies.

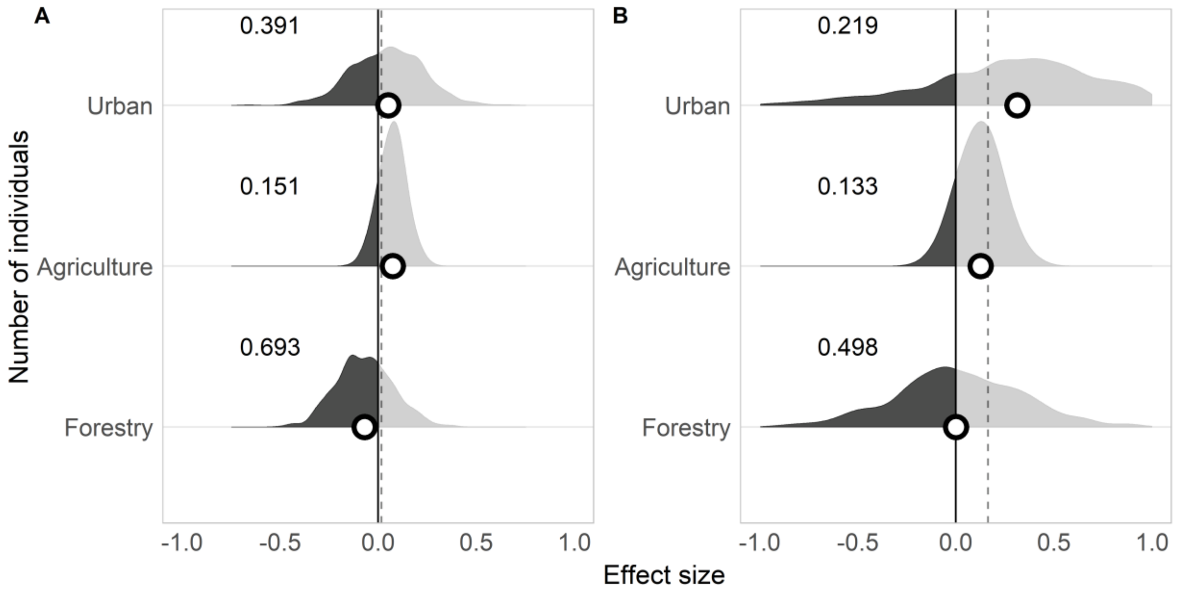


**Figure S3. Density plot of the effect size (log-ratio) between land-use categories at α scale (A), and γ scale (B) of the number of individuals**. The vertical solid black lines show log-ratio equals to 0. The dash vertical lines show the median value of all the log ratios. The numbers show the proportion (out of 1) of log-ratio below 0. The black and gray color show the log-ratio below and above 0.

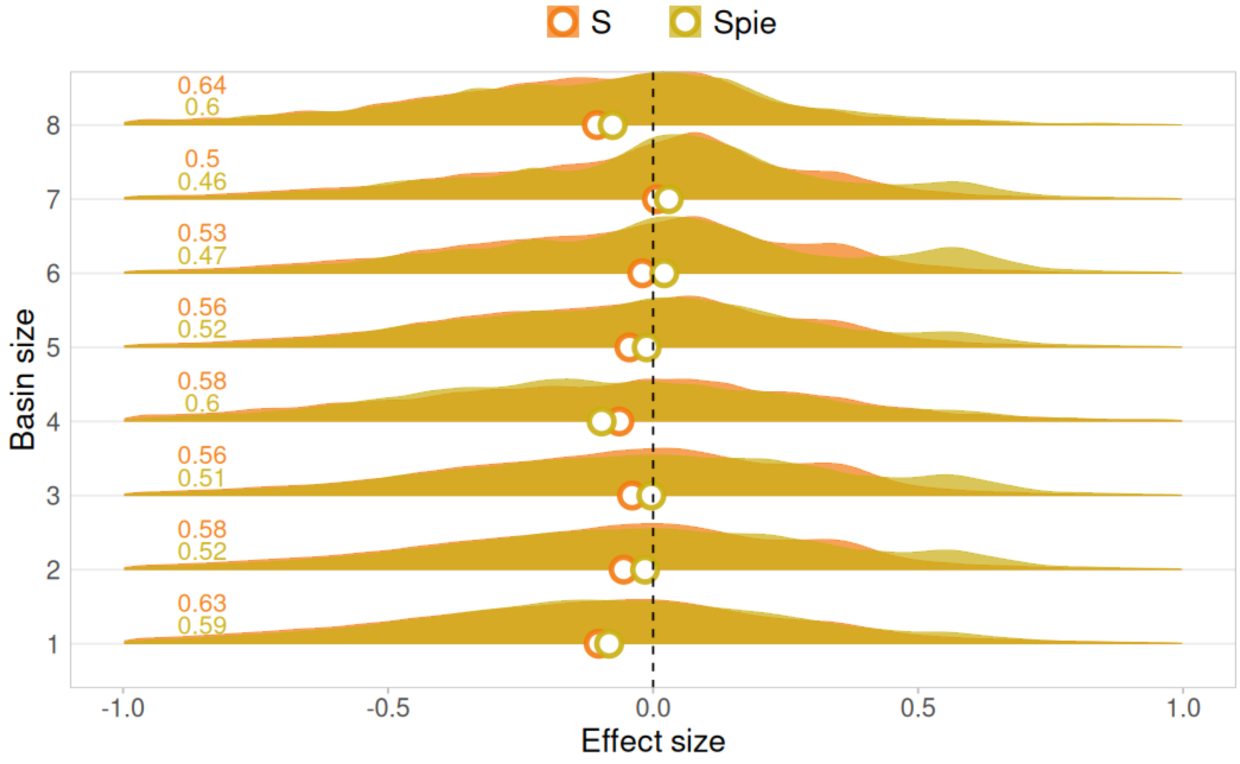

**Figure S4. Density plots of posterior distributions for study-level effect size at α scale grouped by different basin sizes.** The vertical solid black lines show log-ratio equals to 0. The colored dots show the median value of the density plot. The colored numbers show the proportion (out of 1) of log-ratio below 0.

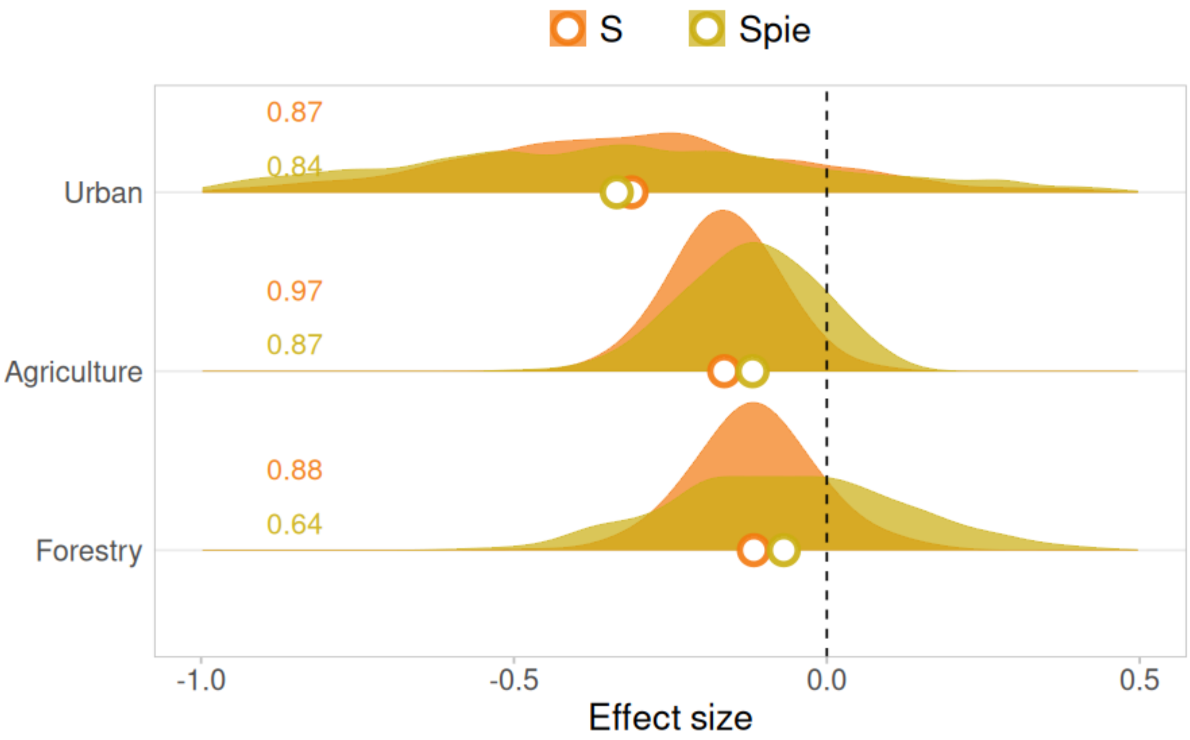


**Figure S5. Density plot of the effect size (log-ratio) between urban, agriculture, forestry and natural vegetation at α scale, using the same datasets as for γ and β diversities.** The vertical solid black lines show log-ratio equals to 0. The colored dots show the median value of the density plot. The colored numbers show the proportion (out of 1) of log-ratio below 0.

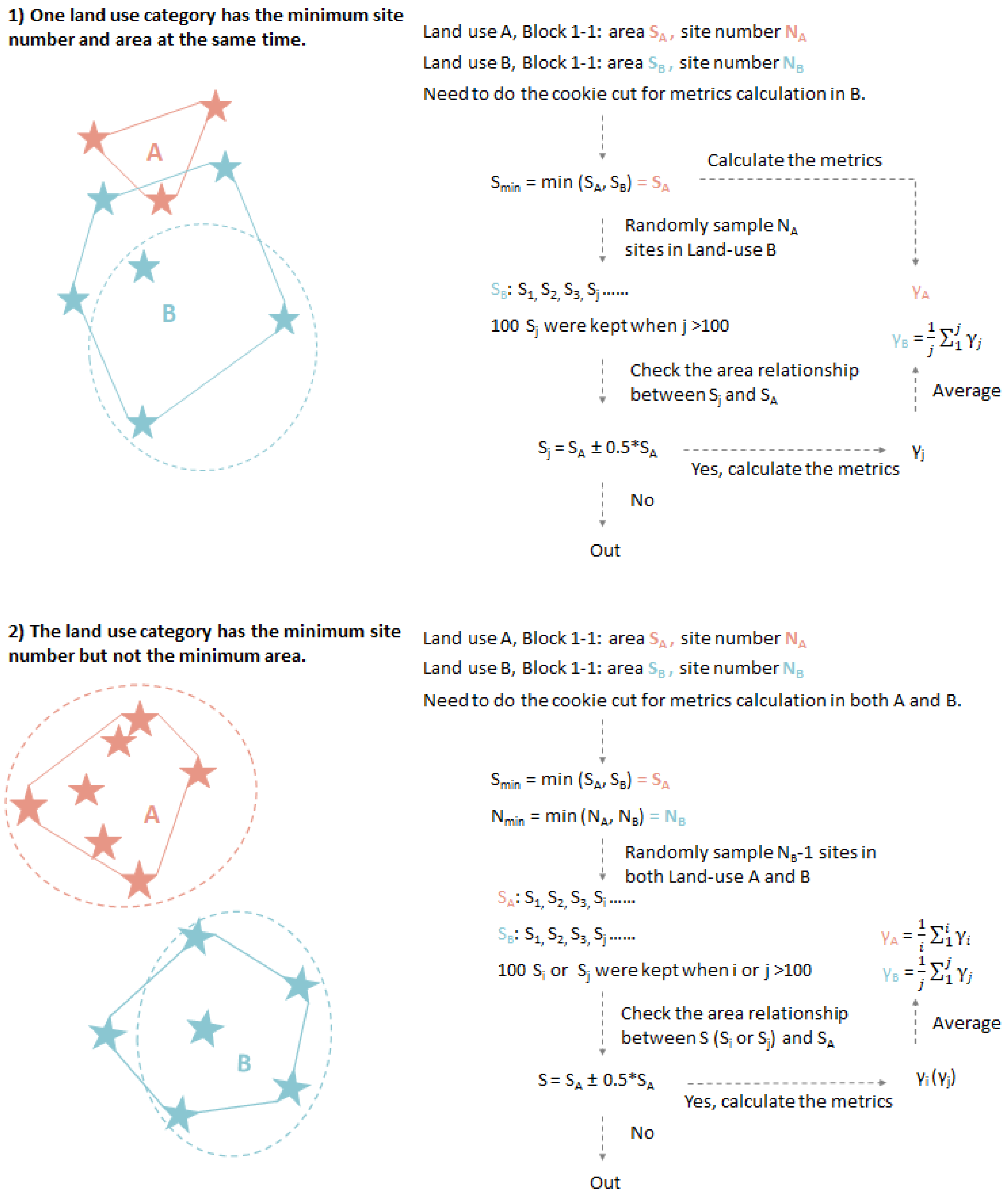


**Figure S6. Workflow for controlling the spatial extent for γ and β diversities under two scenarios**. (1): when comparing land-use A and B, if land-use A had both the smallest number of sites ($N_A$) and the smallest spatial extent ($S_A$), the metrics of land-use A were calculated directly and cookie cutting was applied only to land-use B. We randomly selected 100 combinations of $N_A$ sites from land-use B. For combinations with spatial extents within ±50% of $S_A$, the metrics were calculated, and the mean across these combinations was used as the metric value for land-use B. (2): In cases where land-use A had the smallest spatial extent ($S_A$) but land-use B had the fewest sites ($N_B$), cookie cutting was applied to both land uses. We randomly sampled $N_B$ – 1 sites 100 times in each land use, retaining only combinations with spatial extents within ±50% of $S_A$. The final metric for each land use was the mean of the accepted combinations.

**Supplementary Table 1 Summary of the model testing the overall effect of land use on biodiversity across scales.** For each parameter, the mean estimate, the standard deviation (sd), and the 95% credible interval (Q2.5 and Q97.5) were shown. Rhat is the Gelman-Rubin convergence diagnostic; Bulk- and Tail-ESS are the number of independent samples.

| Metrics | Land use | Estimate | Est.Error | l-95% CI | u-95% CI | Rhat | Bulk_ESS | Tail_ESS |
|---|---|---|---|---|---|---|---|---|
| α_S | Natural vegetation | 2.814725 | 0.068408 | 2.675309 | 2.946717 | 1.010674 | 415.7042 | 797.1325 |
| | Forestry | 2.740425 | 0.088096 | 2.56762 | 2.915213 | 1.002897 | 807.123 | 1883.292 |
| | Agriculture | 2.726491 | 0.067592 | 2.592075 | 2.857291 | 1.008603 | 554.2331 | 1396.413 |
| | Urban | 2.538813 | 0.097795 | 2.344698 | 2.730639 | 1.004907 | 1081.018 | 2000.848 |
| α_Spie | Natural vegetation | 1.602143 | 0.048119 | 1.505926 | 1.696903 | 1.001055 | 3144.962 | 4590.699 |
| | Forestry | 1.555918 | 0.077599 | 1.405978 | 1.711902 | 1.000559 | 4613.265 | 5524.707 |
| | Agriculture | 1.485483 | 0.047878 | 1.393568 | 1.580659 | 1.001814 | 2116.58 | 3622.739 |
| | Urban | 1.283677 | 0.082401 | 1.126634 | 1.445349 | 1.000163 | 4723.426 | 5504.367 |
| γ_S | Natural vegetation | 3.763135 | 0.139489 | 3.495563 | 4.041621 | 1.003531 | 847.504 | 1930.581 |
| | Forestry | 3.699342 | 0.153477 | 3.40165 | 4.0081 | 1.001852 | 988.5962 | 2244.211 |
| | Agriculture | 3.618838 | 0.15328 | 3.320944 | 3.920113 | 1.002847 | 1002.897 | 2490.213 |
| | Urban | 3.488411 | 0.280868 | 2.932558 | 4.04946 | 1.000824 | 2944.79 | 5155.22 |
| γ_Spie | Natural vegetation | 2.157339 | 0.10067 | 1.961228 | 2.35778 | 1.000999 | 1898.452 | 3526.471 |
| | Forestry | 2.098254 | 0.159687 | 1.788394 | 2.41906 | 1.000261 | 2672.39 | 4301.308 |
| | Agriculture | 1.999046 | 0.118606 | 1.76475 | 2.231703 | 1.001388 | 2717.704 | 4460.187 |
| | Urban | 1.585831 | 0.333028 | 0.921872 | 2.252112 | 1.00012 | 3924.568 | 5757.641 |
| β_S | Natural vegetation | 0.976531 | 0.089795 | 0.795122 | 1.150033 | 1.004122 | 1487.865 | 2416.153 |
| | Forestry | 1.025735 | 0.108775 | 0.813349 | 1.243596 | 1.003246 | 2067.982 | 4084.231 |

| | | | | | | | | |
|---|---|---|---|---|---|---|---|---|
| | Agriculture | 0.994755 | 0.087793 | 0.817194 | 1.165873 | 1.003483 | 1871.018 | 3584.908 |
| | Urban | 1.009579 | 0.114261 | 0.778408 | 1.227726 | 1.001395 | 3899.808 | 6271.75 |
| β_Spie | Natural vegetation | 0.512218 | 0.065575 | 0.385248 | 0.641189 | 1.00303 | 2990.436 | 5001.203 |
| | Forestry | 0.525569 | 0.114983 | 0.300984 | 0.752288 | 1.000745 | 4750.68 | 6958.913 |
| | Agriculture | 0.471806 | 0.06787 | 0.334863 | 0.60342 | 1.001066 | 3672.295 | 5740.461 |
| | Urban | 0.296945 | 0.168497 | -0.04005 | 0.632003 | 1.000783 | 5923.846 | 7244.182 |
| α_N | Natural vegetation | 5.674843 | 0.149201 | 5.388243 | 5.974706 | 1.015679 | 290.2494 | 620.7888 |
| | Forestry | 5.59934 | 0.198334 | 5.220462 | 5.98698 | 1.015391 | 485.5443 | 1488.869 |
| | Agriculture | 5.745325 | 0.154572 | 5.443107 | 6.054217 | 1.014611 | 365.6372 | 753.8879 |
| | Urban | 5.721353 | 0.228938 | 5.269321 | 6.168881 | 1.006747 | 686.6156 | 1687.326 |
| γ _N | Natural vegetation | 7.733613 | 0.26847 | 7.222851 | 8.268628 | 1.003876 | 1190.332 | 2502.732 |
| | Forestry | 7.750253 | 0.436918 | 6.874468 | 8.613278 | 1.002636 | 2200.125 | 5713.422 |
| | Agriculture | 7.857177 | 0.271316 | 7.341978 | 8.398298 | 1.003244 | 1313.064 | 2947.628 |
| | Urban | 8.091727 | 0.556698 | 6.974987 | 9.200197 | 1.000406 | 5609.844 | 7695.423 |

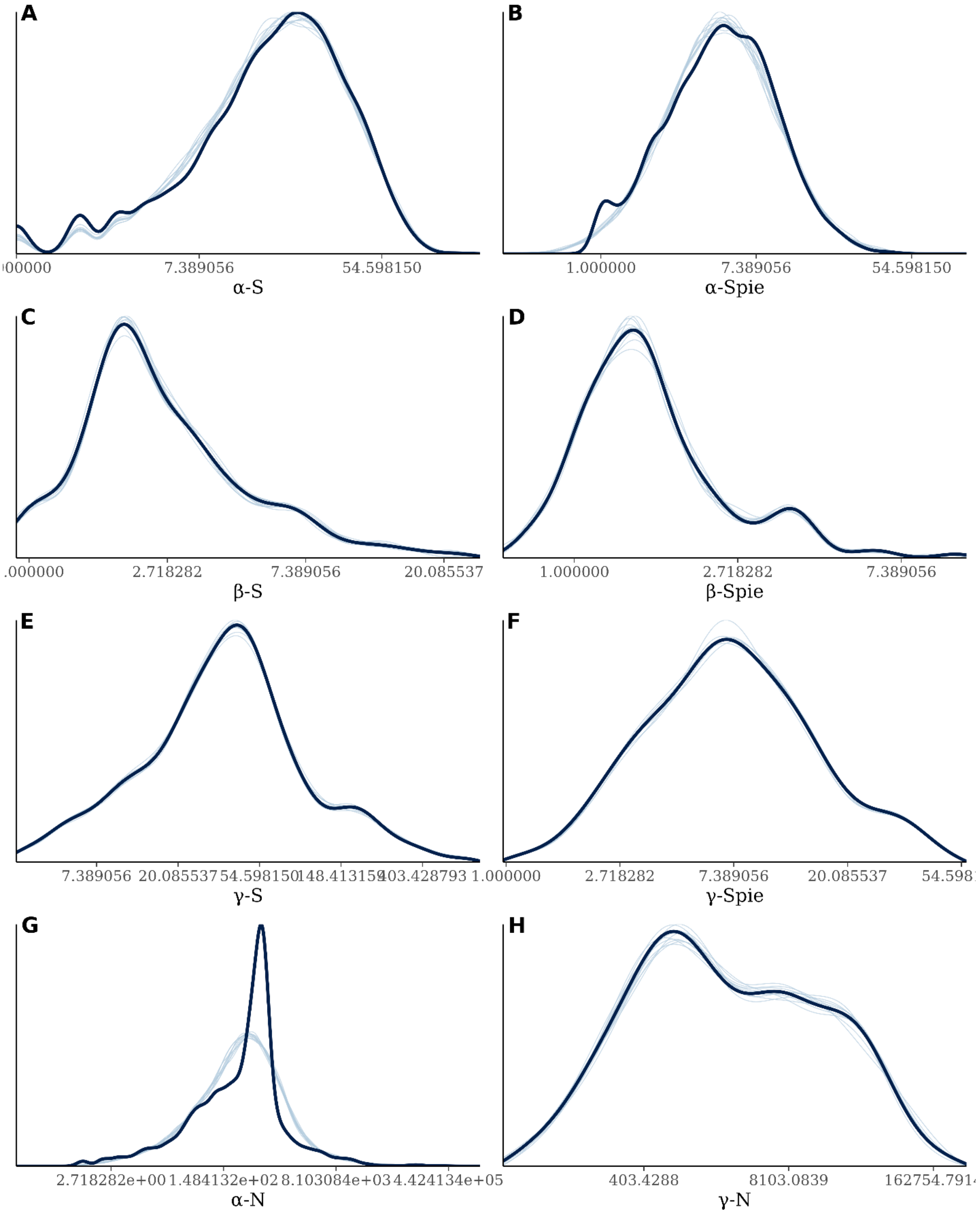


**Figure S7. Comparing kernel density estimates of the metric value (black curves) and predicted value (light blue curves) drawn from the posterior predictive distribution of α-S (A), α-Spie (B), γ-S (C), γ-Spie (D), β-S (E), β-Spie (F), α-N (G), and γ-N (H) models.**

**Supplementary Table 2 Summary of the model testing the overall effect of absolute latitude on land use effect sizes.** For each parameter, the mean estimate, the standard deviation (sd), and the 95% credible interval (Q2.5 and Q97.5) were shown. Rhat is the Gelman-Rubin convergence diagnostic; Bulk- and Tail-ESS are the number of independent samples.

| Metrics | Fixed effect terms | Estimate | Est.Error | l-95% CI | u-95% CI | Rhat | Bulk_ESS | Tail_ESS |
|---|---|---|---|---|---|---|---|---|
| α_S | Intercept | -0.184600 | 0.024727 | -0.233096 | -0.135288 | 1.001159 | 2007.714190 | 2228.570674 |
| | Latitude | 0.001750 | 0.000909 | -0.000022 | 0.003550 | 1.000815 | 2233.741884 | 2735.523358 |
| α_Spie | Intercept | -0.190099 | 0.017370 | -0.224206 | -0.155577 | 1.000267 | 4363.273135 | 3511.656852 |
| | Latitude | 0.001315 | 0.000630 | 0.000104 | 0.002574 | 1.000426 | 4248.151074 | 3245.957635 |
| γ_S | Intercept | -0.163541 | 0.039896 | -0.241094 | -0.085477 | 1.000154 | 3938.700539 | 3387.286914 |
| | Latitude | 0.000091 | 0.001339 | -0.002572 | 0.002777 | 1.001044 | 4178.727607 | 2788.918937 |
| γ_Spie | Intercept | -0.291745 | 0.062061 | -0.414057 | -0.164065 | 1.001395 | 3929.369193 | 2394.854920 |
| | Latitude | 0.001147 | 0.002056 | -0.002910 | 0.005255 | 1.000825 | 4439.158914 | 2694.912954 |
| β_S | Intercept | 0.140724 | 0.033050 | 0.075922 | 0.206186 | 1.001307 | 2537.946842 | 3130.054577 |
| | Latitude | -0.004178 | 0.001114 | -0.006392 | -0.002012 | 1.002521 | 2644.986733 | 2934.230760 |
| β_Spie | Intercept | -0.004026 | 0.040925 | -0.085889 | 0.079303 | 1.001746 | 1513.872368 | 2014.655668 |
| | Latitude | -0.002937 | 0.001388 | -0.005592 | -0.000170 | 1.001376 | 1903.887102 | 2689.053956 |
